# Structural insights and engineering of deep-sea halophilic PET hydrolytic enzymes

**DOI:** 10.1101/2025.08.30.673199

**Authors:** Guoqiang Zhang, Xian Li, Wei Xia, Chengsong Zhang, Shanmin Zheng, Jieke Du, Ning Wang, Xinzhe Chen, Ge Lv, Yushuo Zhao, Tianheng Wang, Yunjun Pan, Mingzhu Zhang, Jian-Wen Huang, Chun-Chi Chen, Siqi Huang, Cheng Zeng, Zhengquan Gao, Jianwei Chen, Guangyi Fan, Xingwang Zhang, Hongliang Wang, Yongfu Sun, Jing Wu, Kun Liu, Rey-Ting Guo, Shengying Li

## Abstract

Pervasive use of polyethylene terephthalate (PET) poses tremendous challenges for global waste management and environmental sustainability, fueling growing interests in enzymatic degradation as an eco-friendly solution. While PET hydrolases hold significant promise, their industrial deployment is hindered by insufficient performance, particularly under high-salinity conditions raised from high substrate loads. Building on our previous discovery of three deep-sea PET hydrolases (dsPETase01, dsPETase05 and dsPETase06) with exceptional halophilicity and PET-degrading activity, we here present their three-dimensional structures and mechanistic characterization. Structural comparison, site-directed mutagenesis, and domain swapping reveal key structural features and the essential role of *C*-terminal acidic residues in salt tolerance. Integrating semi-rational protein design, Transformer-based modeling, and disulfide bond engineering, we synergistically enhance the thermostability and catalytic efficiency of dsPETase05. These findings elucidate the unique structural features and salt adaptation mechanisms of deep-sea halophilic PET hydrolases, and inform the future engineering of biocatalysts for application in harsh industrial environments.

## Introduction

Polyethylene terephthalate (PET) is among the most widely used synthetic polymers worldwide, with an estimated consumption of over 65 million tons in 2021, primarily driven by its broad application in packaging and textiles^1^. However, PET’s chemical stability and resistance to degradation have led to its massive accumulation in global environments. It is estimated that 150–200 million tons of plastic wastes are currently polluting landfills and natural ecosystems^2^, where PET represents one of the dominant polymer pollutants. Therefore, developing sustainable solutions for PET waste management is imperative, and enzymatic degradation has emerged as a highly promising approach^3,4^.

Enzymatic depolymerization offers significant advantages over traditional chemical and mechanical recycling. Specifically, PET hydrolases (PETases) are able to selectively hydrolyze PET into monomers (Supplementary Fig. 1), which can be repolymerized into new PET or repurposed into other value-added products^2,5,6^. The discovery of *Is*PETase from *Ideonella sakaiensis* marked a breakthrough in this field, demonstrating PET degradation at ambient temperatures^7^. However, naturally occurring PETases generally exhibit limited thermostability and reduced activity on crystalline PET, which has greatly restricted their industrial application^8,9^. To address this challenge, the discovery of novel PETases from nature, coupled with advances in enzyme engineering approaches such as (semi-)rational protein design, directed evolution, and high-throughput screening have remarkably enhanced the stability, activity, and substrate specificity of PET hydrolases^2,3,10–12^. Concurrently, the integration of artificial intelligence (leveraging models such as 3D-CNN and Transformer) and landscape profiling further paves the way for more efficient, cost-effective, and scalable PET recycling processes^5,13–15^.

**Figure 1.**
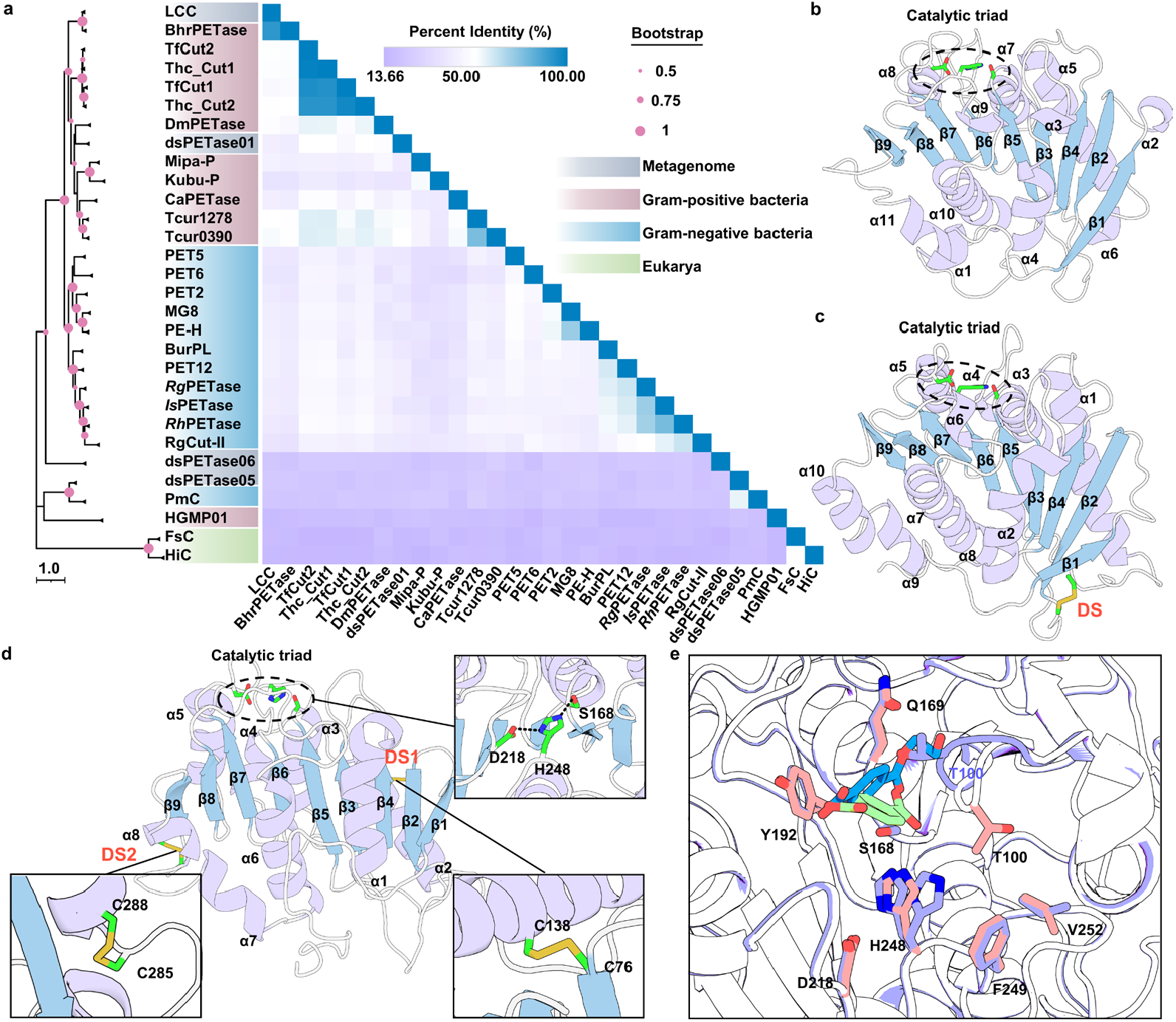
Phylogenetic analysis and overall structures of the three deep-sea halophilic PET hydrolases. **a**, Maximum likelihood phylogenetic tree and percentage identity matrix of the 30 reported PETase sequences. Bootstrap values for 1000 replications are shown as pink circles. Pairwise identity and similarity of the proteins were calculated using the Clustal omega method, and the sequence identity matrix is created using GraphPad Prism 9.0.0 software. **b**, The canonical α/β- hydrolase fold of dsPETase01 (PDB ID #: 9VR5). **c**, Overall structure of dsPETase06_C23A (PDB ID #: 9VR9) with the only disulfide bond shown as sticks in yellow. **d**, Overall structure of dsPETase05 (PDB ID #: 9VR6) with the two disulfide bonds highlighted as sticks in yellow.

Recent major progresses in PET biodegradation have largely been driven by the discovery of novel enzymes from diverse natural environments, with over 120 PET hydrolases experimentally identified to date^16^. Enzymes from extreme habitats, such as composts and glaciers^17–19^, have shown the unique advantages, as wild-type enzymes from these environments often retain high activity under harsh conditions. Deep-sea ecosystems, characterized by high salinity, low temperature, and elevated pressure, represent an untapped reservoir of microbial diversity and enzymatic potential^20^. PET hydrolases have been discovered in deep-sea bacteria^21^ and archaea^22^. Recently, we reported three high-performance PETases derived from deep-sea metagenomes, which require high NaCl concentrations for optimal activity^23,24^, highlighting marine microbiomes as a valuable resource for robust PET hydrolase discovery.

In this study, we resolved the high-resolution crystal structures of three deep-sea halophilic PET hydrolases (dsPETases). Comparative sequence and structural analyses revealed that all three enzymes share a canonical α/β-hydrolase fold and conserved catalytic motifs. The optimal dsPETase05 exhibits three distinctive structural features compared with previously characterized PET hydrolases: (1) several key residues surrounding the catalytic site are markedly different; (2) an additional extended loop is present near the active site; and (3) the *C*-terminal region contains a higher proportion of acidic amino acids. Through site-directed mutagenesis, domain swapping, and functional assays, we validated the functional significance of each feature. Based on these insights, we engineered dsPETase05 by integrating semi-rational protein design, Transformer-based modeling, and strategic disulfide bond incorporation, yielding variants with enhanced thermostability and PET degradation efficiency. These findings offer structural and mechanistic insights into halophilic PET hydrolases and provide a rational basis for developing more robust biocatalysts for PET waste management.

## Results

### Phylogenetic analysis of PET hydrolases

PETases are a group of enzymes belonging to the α/β hydrolase superfamily, characterized by highly conserved structural folds despite low sequence identity^25,26^. To understand the evolutionary relationships among the three deep-sea halophilic PETases we discovered recently^23^, we built a phylogenetic tree comprising dsPETase01, dsPETase05, dsPETase06, alongside 27 previously characterized PET hydrolases (Fig. 1a). The 30 enzymes form three distinct clades: (1) Gram-positive bacterial hydrolases dominated by thermophilic enzymes (*e*.*g*., LCC and TfCut2), (2) Gram-negative bacterial hydrolases represented by mesophilic counterparts such as *Is*PETase, and (3) fungal hydrolases exemplified by FsC and HiC.

The three deep-sea PETases exhibit divergent phylogenetic distributions. Specifically, dsPETase01 is clustered within the Gram-positive bacterial clade, while dsPETase05 and dsPETase06 show closer phylogenetic distance to fungal PET hydrolases and lack clear taxonomic affiliation with either Gram-positive or Gram-negative bacterial groups. The most active dsPETase05 shows the closest evolutionary proximity to PmC, a Gram-negative bacterial hydrolase, whereas dsPETase06 occupies a distinct, phylogenetically independent position in the tree. These findings reveal substantial sequence divergence and suggest distinct evolutionary trajectories for the three deep-sea PETases.

### Overall structures of deep-sea halophilic PETases

To elucidate structural basis underlying the unique catalytic properties of the three deep-sea halophilic PETases, we determined their high-resolution crystal structures (Supplementary Table 1). All three enzymes adopt the canonical α/β-hydrolase fold, yet display several distinctive structural features. The structure of dsPETase01 was solved at 1.7 Å resolution in space group *P*2_1_ (PDB ID #: 9VR5), with two chains observed in an asymmetric unit (Fig. 1b, Supplementary Fig. 2). The structure comprises a central twisted β-sheet composed of nine β-strands flanked on both sides by eleven α-helices. The conserved catalytic triad residues S159, H237, and D205 are consistent with other PET hydrolases^8,9^. According to established classification schemes^8^, dsPETase01 is likely a Type I enzyme, similar to LCC. Intriguingly, dsPETase01 lacks any disulfide bond, distinguishing it from most characterized PET hydrolases.

For dsPETase06, initial crystallization attempts of the wild-type protein was unsuccessful, potentially due to structural heterogeneity introduced by Cys23 at the *N*- terminus. Subsequent expression of the C23A mutant yielded high-resolution crystals diffracting to 1.38 Å, though residues 23-34 at the *N*-terminus remained unresolved (PDB ID #: 9VR9). The refined structure comprises ten α-helices and a 9-stranded β- sheet core. Within the catalytic triad (Ser156-His234-Asp205), Ser156 acts as the nucleophile targeting the carbonyl carbon of the scissile ester bond. A single intramolecular disulfide bond (C47-C67) was identified to locate proximal to the *N*- terminal domain (Fig. 1c).

Residues of catalytic triad and oxyanion hole are shown as sticks on the top right. **e**, Superposition of dsPETase05 in complex with TPA (green, PDB ID #: 9VR7) and dsPETase05_S168A in complex with MHET (blue, PDB ID #: 9VR8). Catalytic triad and oxyanion hole residues (purple for 9vR7 and salmon for 9VR8) are shown as sticks. In **b**−**d**, the numbered β-sheets, α-helices, and loop regions are colored differently for clarity; amino acids that constitute the catalytic triad are shown in sticks in black dash line circle.

Structural characterization of dsPETase05, the most catalytically efficient among the three halophilic PETases, revealed distinctive molecular features. The crystal structure, determined at 2.06 Å resolution (space group *I*2_1_3), contains a single molecule in the asymmetric unit (PDB ID #: 9VR6). The enzyme adopts a canonical α/β-hydrolase fold comprising nine β-strands flanked by eight α-helices. A conserved catalytic triad (S168-H248-D218) coordinates nucleophilic attack on the substrate ester bond. Two intramolecular disulfide bonds exist: DS1 (C76-C138) stabilizes the *N*-terminal domain, while DS2 (C285-C288) reinforces the *C*-terminal region (Fig. 1d).

We next compared the surface hydrophobicity and charge distribution of the three halophilic dsPETases with *Is*PETase and LCC (Supplementary Fig. 3). While all five enzymes possess similar hydrophobic pockets, their global and local surface charge distributions differ markedly. Such disparities are likely to influence substrate recognition and catalytic efficiency. In addition, structural superposition analysis revealed that dsPETase05 exhibits the greatest structural divergence among the five proteins, as evidenced by the root mean square deviation (RMSD) values when aligned with the other PETases (Supplementary Fig. 4).

To probe enzyme-substrate interactions, we also obtained high-resolution complex structures by co-crystallizing wild-type dsPETase05 with terephthalic acid (TPA) (1.5 Å, PDB ID #: 9VR7) and an inactive mutant (S168A) with mono(2-hydroxyethyl) terephthalate (MHET) (1.49 Å, PDB ID #: 9VR8). Comparative analysis revealed minimal structural perturbation (Cα RMSD = 0.096 Å) between these two complexes (Fig. 1e). Notably, the binding positions of the ligands exhibit subtle differences, and the conformational flexibility in residues T100 and T101 was clearly observed. Upon binding to MHET, these residues adopt distinct orientations. This is corroborated by analysis of the electron density maps (Supplementary Fig. 5). The substrate-binding cleft is delineated by a conserved aromatic-hydrophobic pocket formed by Y192, V252, F249, H167, T100, and Q169.

### Structural features of dsPETase05

Structural alignment with the two most-studied PET hydrolases (*i*.*e*., LCC and *Is*PETase) demonstrated that dsPETase05 retains the conserved catalytic triad (S168-D218-H248), while exhibiting structural and sequence divergence at critical functional sites including the oxyanion hole, wobbling pairs, substrate-binding regions, and disulfide bond configurations (Fig. 2a). Comparative sequence and structural analyses across representative PET hydrolases further identified a unique extended loop adjacent to the active site in dsPETase05 (Fig. 2b). We also identified several key amino acid residues exhibiting notable divergence at functionally important sites. These structurally mapped residues were subsequently selected for site-directed mutagenesis and enzymatic activity assays, enabling systematic evaluation of their contributions to the catalytic mechanism (Fig. 2c).

**Figure 2.**
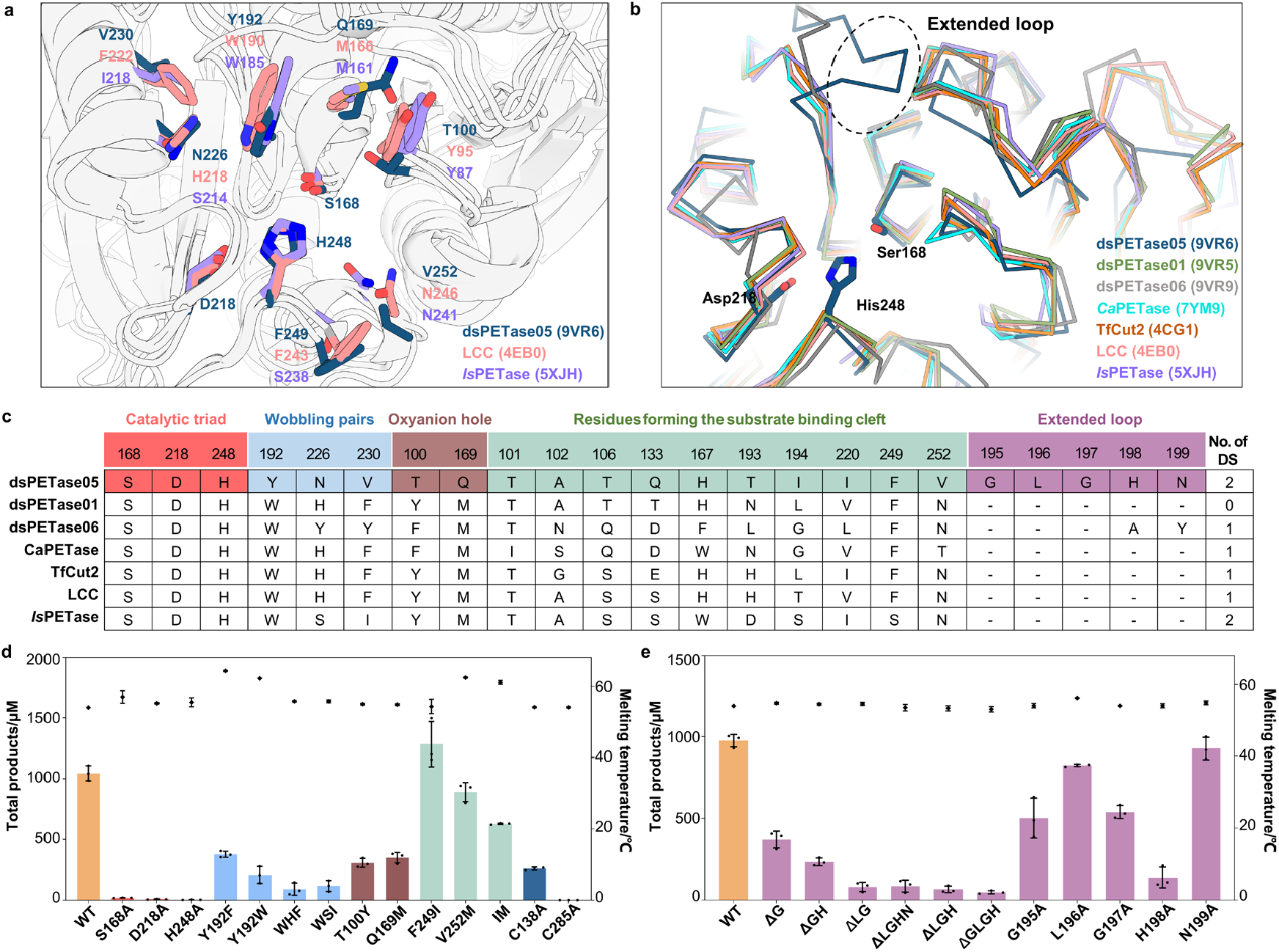
Validation of key structural features in dsPETase05. **a**, Spatial distribution of critical amino acid residues in the structures of dsPETase05, LCC, and *Is*PETase. **b**, Unique extended loop in dsPETase05, with its catalytic triad illustrated in sticks. **c**, Comparative analysis of key residues, extended loop, and the number of disulfide bond(s) across different enzyme variants. **d**, Mutagenesis analysis of critical residues. **e**, Amino acid residue deletion and alanine scanning mutagenesis of the extended loop. Reactions were performed with 50 nM enzyme in 5.3 M NaCl, pH 9.0 Tris-HCl buffer for 24 h. Wild-type dsPETase05 served as the control. Enzymatic activity (product yield, bars) and thermal stability (melting temperature *T*_m_, ◆) are shown.

Alanine substitution of any residue within the catalytic triad (*i*.*e*., S168A, H248A, and D218A) resulted in a near-complete loss of enzymatic activity, underscoring their indispensable roles in catalysis (Fig. 2d). To further elucidate the functional significance of disulfide bonds in dsPETase05, we generated two cysteine mutants, C138A and C285A. Disruption of either disulfide bond significantly diminished the PET-hydrolyzing activity of dsPETase05, with the C285A mutant exhibiting a particularly substantial loss of enzymatic function (Fig. 2d). These results strongly suggest that both disulfide bonds are essential for maintaining the structural integrity of dsPETase05 and preserving its catalytic function.

Structural comparison of the dsPETase05-MHET complex with LCC and *Is*PETase revealed significant divergence in the oxyanion hole. Specifically, dsPETase05 harbors T100 and Q169 at positions corresponding to Y95 and M166 in LCC and Y87 and M161 in *Is*PETase, respectively (Fig. 2a). Additionally, we identified a distinctive feature in dsPETase05: a tyrosine residue (Y192) replaces the tryptophan conserved at the equivalent position in both LCC (W190) and *Is*PETase (W185), a residue previously reported to play a critical role in PET hydrolysis^3,9,27^.

To evaluate the functional impact of these differences, we performed site-directed mutagenesis analysis. As shown in Fig. 2d, the T100Y and Q169M mutants exhibited significantly reduced enzymatic activity, retaining only 29.1% and 33.2% of the activity of wild-type enzyme, respectively. Similarly, the Y192F and Y192W mutations led to substantial activity loss, preserving merely 35.9% and 19.8% of the original activity. Combinatorial mutations (Y192W/N226S/V230I, WSI; Y192W/N226H/V230F, WHF) further reduced activity to 11.2% and 8.7% of the wild-type level, respectively. These results collectively demonstrate that dsPETase05 possesses a unique oxyanion hole architecture and tyrosine residue configuration, distinguishing it from other well-characterized PET hydrolases including LCC^19^, *Is*PETase^7^, CaPETase^28^, and TfCut2^29^.

Following the successful engineering strategy of LCC-ICCG^2^, we introduced targeted mutations F249I, V252M, and combinatorial mutant IM (F249I/V252M) at positions 249 and 252 within the substrate-binding pocket of dsPETase05. Our enzymatic assays demonstrated that among these mutants, only F249I exhibited enhanced activity, displaying 1.23-fold higher catalytic efficiency compared to the wild-type enzyme (Fig. 2d). This positions F249I as a strategic candidate for combinatorial mutagenesis in further enzyme engineering endeavors.

To investigate the functional relevance of the extended loop in dsPETase05, we performed both amino acid deletion and alanine scanning mutagenesis. Sequential deletion of residues from this loop resulted in a progressive decline in enzymatic activity under high-salt conditions (Fig. 2e), indicating that this loop is crucial for maintaining enzyme structure and/or catalytic function. Alanine scanning further identified H198 as essential for enzymatic activity. These findings were corroborated by the result that the H198 deletion mutants exhibited severely impaired activity.

### *C*-terminal acidic amino acids are crucial for the halophilic property of dsPETase05

A commonly observed feature of halophilic enzymes is the enrichment of acidic amino acids (aspartate and glutamate) on their protein surfaces, which has been proposed to enhance protein solubility and stability under high-salt conditions^30,31^. Sequence and structure analysis of dsPETase05 revealed a total of 18 acidic residues, 10 of which are located in the *C*-terminal region, specifically after the last catalytic triad residue H248 (Fig. 3a). In contrast, *Is*PETase, which we have previously demonstrated to be non- halotolorant^23^, contains only 12 acidic amino acids, with merely 5 located at its *C*- terminal region following H237 (Fig. 3b).

**Figure 3.**
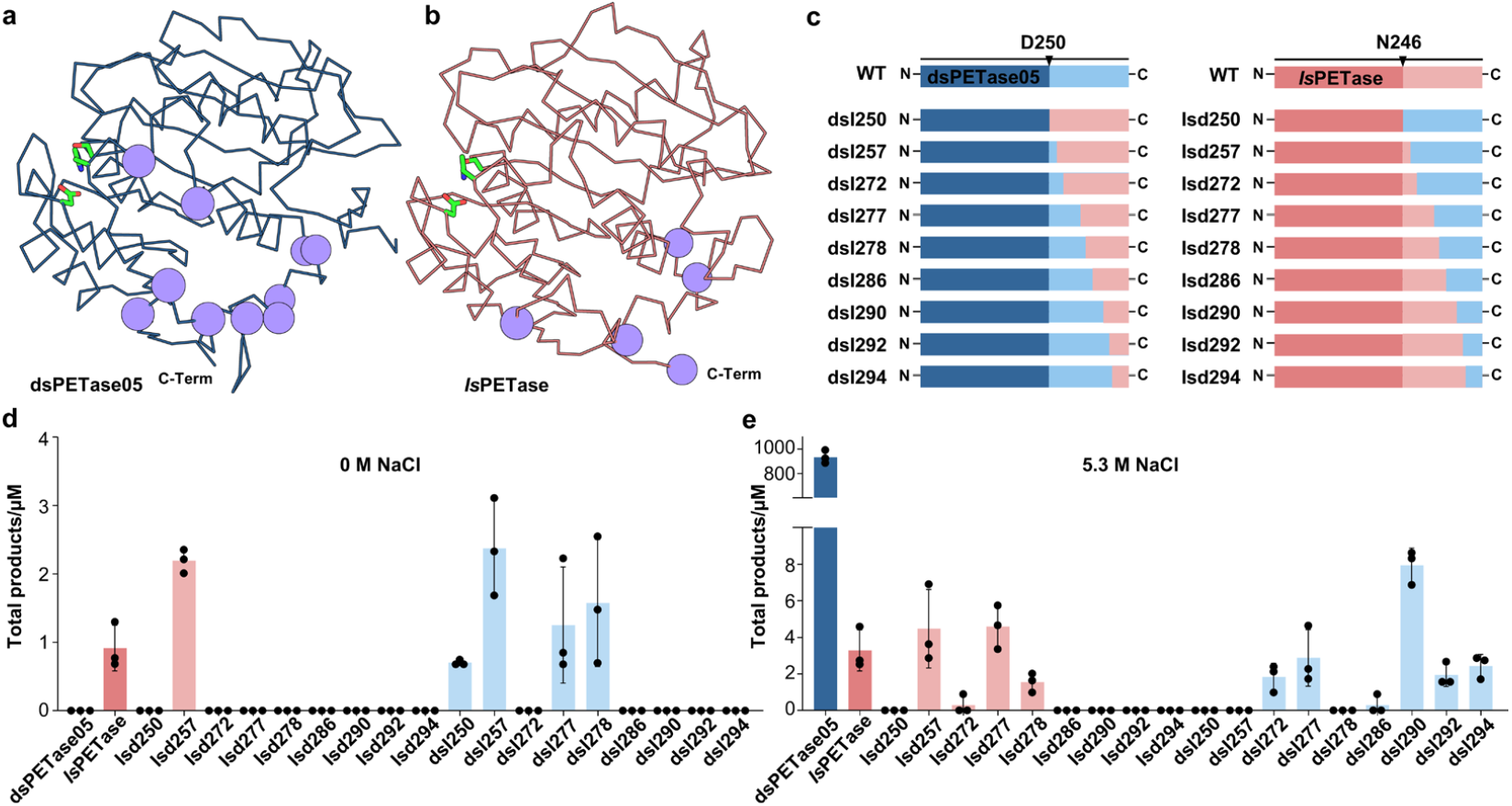
Exploring the halophilic mechanism of dsPETase05. **a**, Distribution of *C*-terminal acidic residues in dsPETase05: protein backbone depicted with dark blue lines, purple spheres representing acidic residues, and catalytic triad shown as sticks; **b**, Distribution of *C*-terminal acidic residues in *Is*PETase: dark pink lines for protein backbone, purple spheres for acidic residues, and sticks for catalytic triad; **c**, Schematic exchange of *C*-terminal regions in different lengths within the terminal hydrophilic domains of dsPETase05 and *Is*PETase (WT dsPETase05: dark blue bar - main body, light blue bar - acidic *C*-terminus; WT *Is*PETase: dark pink bar - main body, light pink bar - *C*-terminus); **d** and **e**, Enzymatic activities of *C*-terminal domain-swapped mutants under 0 M NaCl (**d**) and 5.3 M NaCl (**e**) conditions. Reactions were performed with 50 nM enzyme in pH 9.0 Tris-HCl buffer for 24 h. Column heights indicate the means of triplicate experiments, circles represent individual data points, and error bars denote standard deviations.

To evaluate the functional significance of the *C*-terminal acidic residues, we adopted a domain swapping strategy to construct a series of chimeric enzymes. Specifically, we replaced the *C*-terminal segments of dsPETase05 and *Is*PETase with the corresponding regions from each other, thereby generating 18 chimeric mutants (9 based on the dsPETase05 scaffold and 9 on the *Is*PETase scaffold; Fig. 3c). Notably, substitution of the *C*-terminal region of dsPETase05 with that of *Is*PETase—which is relatively depleted in acidic residues—resulted in chimeric enzymes exhibiting significantly enhanced catalytic activity under salt-free conditions compared to wild-type dsPETase05 (Fig. 3d). In contrast, transplantation of the acidic residue-rich *C*-terminal regions from dsPETase05 into *Is*PETase substantially improved halotolerance of the resulting chimeric enzymes (Isd257 and Isd277), as evidenced by their substantially increased enzymatic activity at 5.3 M NaCl relative to native *Is*PETase. These findings indicate that *C*-terminal acidic amino acids play a critical role in determining the halophilic properties of dsPETase05.

To further delineate the specific acidic residues responsible for halotolerance, we conducted site-directed mutagenesis on dsPETase05. Six mutants (dsIr1–dsIr6) were generated by individually substituting each targeted acidic residue in the *C*-terminal region of dsPETase05 with the corresponding amino acid found in *Is*PETase (see Supplementary Table 2 for details). Under 5.3 M NaCl, dsIr6 exhibited sharply decreased activity compared to dsPETase05 and other mutants (Supplementary Fig. 6a), indicating that this specific residue is critical for maintaining halotolerance. In contrast, under salt-free conditions, the dsIr5 mutant displayed increased activity relative to dsPETase05, other mutants, and *Is*PETase (Supplementary Fig. 6b). These results highlight the essential roles of D237 and E277 in the *C*-terminal region in maintaining the halophilic adaptation and catalytic performance of dsPETase05.

### Rational design and directed evolution of dsPETase05 for enhanced PET hydrolysis

The PET-hydrolyzing activity observed in PETases is widely regarded as a promiscuous function that has been selectively enhanced by environmental pressures to become a primary catalytic role^32^. To date, many highly active PET-degrading biocatalysts reported are thermotolerant enzymes exhibiting optimal PET hydrolysis near the PET glass transition temperature (65–81 °C)^33,34^. Thus, to enable industrial application of PET hydrolases for PET depolymerization and recycling, protein engineering is typically required to enhance their kinetic and thermal stability as well as catalytic efficiency. Accordingly, we implemented three strategies in order to improve the PET-degrading capacity of dsPETase05: (1) Machine learning-guided mutational site identification: a Transformer model was employed to predict the probability of amino acid variation from the evolutionary landscape^13^, followed by high-throughput fluorescence screening^12^ to rapidly validate activity-enhancing mutations (Fig. 4a); (2) Single-site saturation mutagenesis: starting from the F249I mutant with enhanced activity, we performed comprehensive mutagenesis at this position to identify superior amino acid replacements (Fig. 4b); (3) Disulfide bond engineering: using systematic homology-driven disulfide bond engineering strategy^35^, we predicted viable mutation pairs within 7.5 Å in the dsPETase05 structure for introducing disulfide bonds (Fig. 4c and Supplementary Table 3).

**Figure 4.**
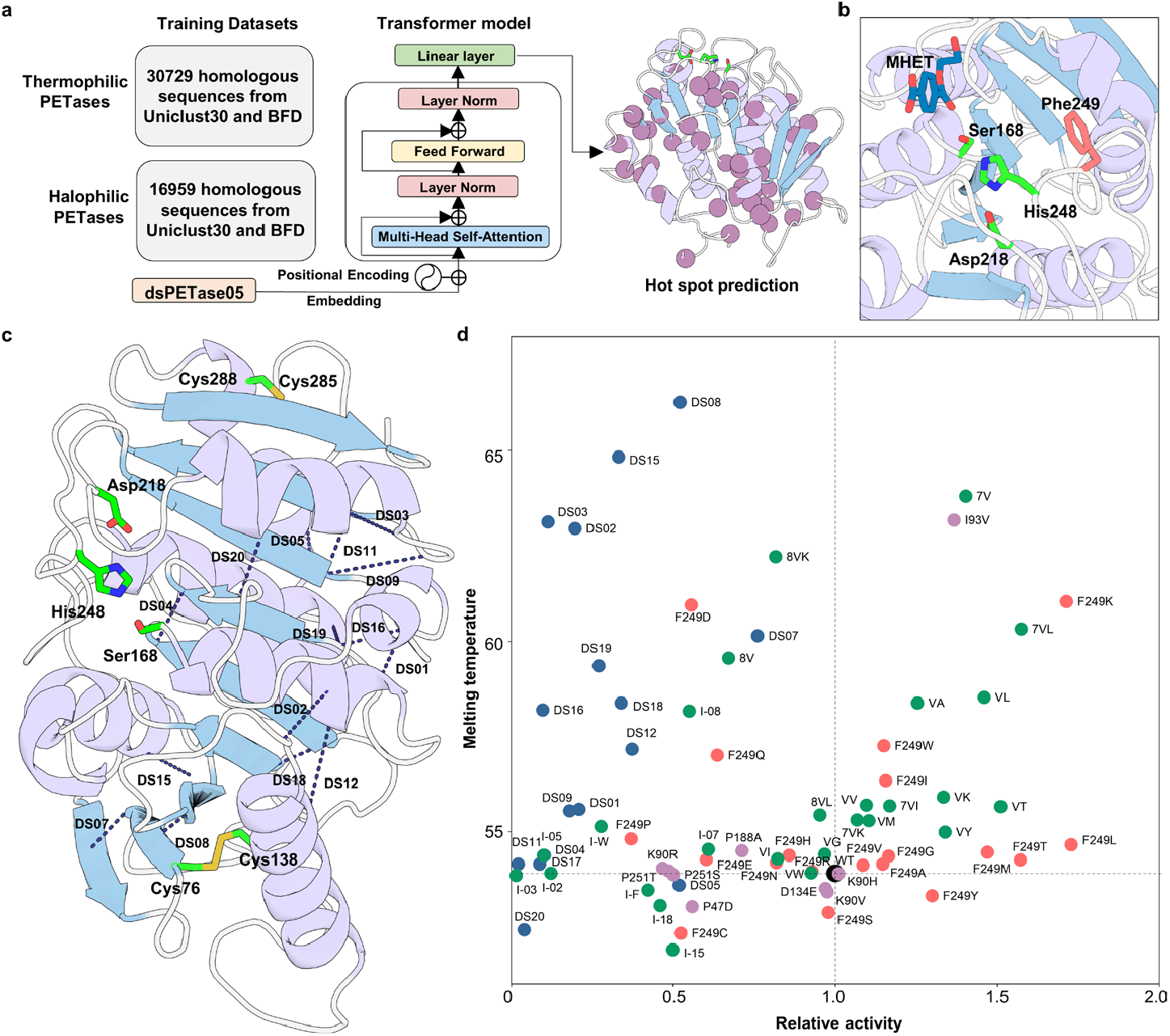
Activity and thermostability optimization of dsPETase05 by enzyme engineering. **a**, Prediction of potentially beneficial mutations using the Transformer model, with 45 candidate amino acid positions (highlighted by dark purple spheres) selected through computational screening. **b**, Structural analysis of the dsPETase05-MHET complex, illustrating residue F249 in salmon, the MHET moiety in marine, and catalytic triad in green. **c**, Disulfide bond engineering: amino acid pairs within 7.5 Å were selected for covalent linkage design, with inter-residue distances indicated by dark blue dashed lines. **d**, Thermodynamic characterization and catalytic performance: thermal stability (*T*_m_) and relative PET degradation activity of dsPETase05 variants. Data derived from an independent experimental trial. Mutation types are color-coded: wild type (black), Transformer-predicted substitutions (lilac), F249 saturation mutagenesis (salmon), disulfide bond-engineered variants (dark blue), and combinatorial mutants (green).

First, the Transformer-based machine learning model prediction yielded 45 candidate residues with potential to improve catalytic activity. For each predicted site, an NNK-based site-saturation mutagenesis library was constructed, with 96 mutants per 96-well plate to ensure approximately 95% coverage at each position, resulting in a total of ∼4,500 variants^36^. Subsequently, we employed a dual fluorescence-based high-throughput screening (HTS) assay previously developed by our group^12^, which enables rapid and quantitative assessment of PET hydrolysis products in 96-well plates. The top 20 performing variants were subjected to sequencing, and after removing wild-type and redundant sequences, nine unique mutant proteins were identified. Subsequent protein purification and enzymatic assays revealed that, as shown by the lilac markers in Fig. 4d, only the I93V variant exhibited simultaneous improvements in both enzymatic activity and melting temperature (*T*_m_), while other mutants demonstrated comparable or inferior performance relative to dsPETase05. Consequently, I93V was prioritized as the combinatorial mutagenesis target for subsequent optimization.

In dsPETase05, F249 is located adjacent to the histidine residue of the catalytic triad. F249I mutant was found to enhance enzymatic activity, supporting the hypothesis that reducing steric hindrance at this position may facilitate substrate access to the active site and hence improve catalytic efficiency^2^. Thus, we performed single-site saturation mutagenesis at F249. Nine mutants showed both higher enzymatic activity and increased *T*_m_ compared to the wild-type enzyme, with F249K displaying the most pronounced improvement (Fig. 4d).

To further improve the thermal stability of dsPETase05, we employed a structure-guided disulfide bond engineering strategy. Drawing inspiration from previous successes—the introduction of disulfide bonds in LCC, which resulted in a 6.8 °C increase in *T*_m_ and substantially improved activity in the LCC-ICCG variant^2^—we designed and introduced 15 disulfide bond mutations into dsPETase05. Enzymatic assays (Fig. 4d, dark blue circular markers) demonstrated that, while most disulfide bond mutants exhibited elevated *T*_m_ values, their catalytic activities were generally lower than that of wild-type dsPETase05.

To balance the trade-off between enhanced thermostability and catalytic activity, we further combined the beneficial single-point mutations (such as I93V and F249L) with selected disulfide bond mutants that exhibited notable increases in *T*_m_, resulting in 28 soluble mutants (Supplementary Table 4). As a result, the 7V mutant (DS07/I93V) achieved a *ΔT*_m_ of 9.9 °C and exhibited 1.4-fold increase in PET hydrolytic activity. Notably, the 7VL variant (DS07/I93V/F249L) maintained a *ΔT*_m_ of 6.4 °C while demonstrating nearly 1.6-fold improvement in PET-specific activity (Fig. 4d). Collectively, these results provide a robust foundation for further optimizing this halophilic enzyme toward more efficient PET degradation and recycling.

## Discussion

Recent breakthroughs in enzymatic PET depolymerization have highlighted the importance of discovering new PET hydrolases from diverse natural habitats, understanding their adaptation mechanisms, and optimizing their industrial properties^2,7,37^. To make PET biodegradation and recycling more economically viable, achieving high substrate loading and low enzyme dosage has become a key direction for industrial application^13^. High PET loading would inevitably lead to the accumulation of high concentrations of the acidic product TPA, which requires the addition of bases such as NaOH for neutralization, resulting in the generation of large amounts of salts at molar levels. The halophilic enzymes we recently discovered from deep-sea environments, such as dsPETase05, exhibit optimal activity at nearly saturated NaCl concentrations^23^, which are even higher than the salt concentrations generated during neutralization in high-substrate PET depolymerization processes. This remarkable salt tolerance suggests that understanding the behind mechanisms and engineering better mutants will be instrumental in overcoming product inhibition and other challenges associated with industrial PET recycling. Therefore, the discovery, mechanistic elucidation, and engineering of halophilic PET hydrolases are of great importance to advance this field.

During the preparation of this manuscript, a new halophilic PET hydrolase was discovered from the Tara Ocean metagenomic database, and structural analysis also revealed the presence of an extended loop^38^, a feature observed in dsPETase05 and also can be found in a previously identified PET-active lipase (PDB ID code: 2FX5). Our in-depth structural and mutational analyses of dsPETase05 characterized this loop and demonstrated its critical role in structural stability under salt stress, and also revealed that the enrichment of surface acidic residues and a distinctive *C*-terminal domain are critical for halotolerance and enzyme stability, which is in line with other halophilic enzymes such as halophilic xylosidase^39^ and malate dehydrogenase^31^. Integration of machine learning-guided mutagenesis, saturation mutagenesis, and disulfide bond engineering achieved synergistic improvements in both thermostability and PET hydrolytic activity for dsPETase05. Notably, our approach aligns with the emerging consensus that multi-dimensional optimization targeting thermostability and activity is essential for the practical deployment of PET hydrolases at scale^10,13,15,34,40,41^.

Given the demonstrated advantages of halophilic PET hydrolases in overcoming challenges during industrial PET depolymerization, further discovery and mechanistic understanding of new enzymes with desirable properties are particularly crucial for advancing sustainable plastic recycling. Marine environments, as a vast and largely untapped reservoir of enzymes, offer significant potential in this regard^23,42^. Recent studies have revealed that microorganisms and enzymes derived from the ocean can degrade plastics including PET and polyethylene^43–45^, and these marine enzymes often exhibit unique adaptations—such as halophilicity, psychrophilicity, or thermophilicity—alongside novel protein structures that may help address current limitations in plastic biodegradation. Therefore, integrating the exploration and engineering of marine enzyme resources with advanced protein design strategies and multidisciplinary collaborations will be essential to develop robust biocatalysts and accelerate the industrial application of enzymatic plastic recycling.

## Materials and Methods

### Materials and reagents

The following materials and reagents were obtained from commercial sources: amorphous PET film (GfPET, ES301445) from Goodfellow (Huntingdon, England); terephthalic acid (TPA), bis(2-hydroxyethyl) terephthalic acid (BHET), mono(2-hydroxyethyl) terephthalic acid (MHET) and 1,1,1,3,3,3-Hexafluoro-2-propanol (HFIP) from Aladdin (Shanghai, China); fluorescein dilaurate (FDL) and BugBuster from Sigma-Aldrich (Darmstadt, Germany); antibiotics and isopropyl-*β*-D-thiogalactopyranoside (IPTG) from Solarbio (Beijing, China); PrimeSTAR Max DNA Polymerase and rTaq DNA Polymerase from Takara (Dalian, China); and ClonExpress II One Step Cloning Kit from Vazyme (Nanjing, China).

### Plasmid construction

The encoding genes for dsPETase01, dsPETase05 and dsPETase06 with signal peptide truncation were chemically synthesized by BGI (Shenzhen, China). All plasmids were constructed by seamless cloning using ClonExpress II One Step Cloning Kit. For crystallographic studies, these genes were sub-cloned into pET32a, whereas for enzymatic characterization, both wild-type and mutant variants were sub-cloned into pET28a. The variants were generated through PCR-based site-directed mutagenesis. The constructed plasmids and their respective mutagenesis primers used in this study are shown in Supplementary Table 5. All constructed plasmids were validated by DNA sequencing.

### Recombinant protein expression and purification

The constructed plasmids were transformed into *E. coli* BL21(DE3) cells and grown in Luria–Bertani medium containing 100 mg L^-1^ ampicillin (when using pET32a) or 50 mg L^-1^ kanamycin at 37 ℃, 220 rpm. When optical density at 600 nm (OD_600_) reached 0.8, 0.3 mM IPTG was added to induce protein expression at 16 ℃ for 18 h. Cell pellets were collected via centrifugation at 5,000 *g* for 8 min and then resuspended in lysis buffer (25 mM Tris-HCl, 150 mM NaCl and 20 mM imidazole, pH 7.5) and then disrupted using a French press machine. Cell lysates were centrifuged at 17,000 *g* and 4 ℃ for 1 h, and the resulting supernatant was loaded onto a Ni-NTA (Sangon Biotech, Shanghai, China) column, the target protein was eluted with a 20-300 mM linear gradient of imidazole. Fractions containing target protein were collected and dialyzed against a buffer containing 25 mM Tris-HCl (pH 7.5), and 150 mM NaCl. Proteins expressed from pET28a were purified directly as described above for activity assays, whereas pET32a-derived proteins underwent additional steps for Trx tag removal and purification: the Trx tag was cleaved by Tobacco etch virus protease, followed by second Ni-NTA chromatography to collect the untagged proteins. The purified proteins were then concentrated for crystallization.

### Crystallization and structure determination

All protein crystallization trials were conducted using the sitting drop vapor-diffusion method at 25 ℃ by mixing 1 μL protein solution and 1 μL reservoir solution in 96-well Crys chem plates and equilibrated against 100 μL of the reservoir solution. The growing conditions for crystals with the best diffraction quality were as follows: dsPETase01 (25% PEG3350, 0.1 M sodium fluoride); dsPETase05 (2.8 M sodium acetate trihydrate pH 7.0, 0.1 M Bis-Tris propane, pH 7.0); dsPETase06_C23A (30% v/v PEG400, 0.1 M HEPES pH 7.5, 0.2 M magnesium chloride hexahydrate); substrate analogues and products were used for socking or co-crystallization to obtain the complex structures. The X-ray diffraction datasets were collected at beamlines TPS-07A of the National Synchrotron Radiation Research Center (NSRRC) and a Bruker D8 Venture coupled with a CMOS-PHOTON II detector at Hubei University. HKL-2000 (version 722) software and Proteum 3 (Bruker AXS GmbH) were used to process the X-ray diffraction datasets.

### PET hydrolase activity assays

Amorphous PET substrates were prepared by cutting GfPET film (ES301445, Goodfellow) into 6-mm-diameter round pieces. The enzymatic hydrolysis was performed by incubating PET pieces with 50 nM enzyme in 500 μL Tris-HCl buffer (pH 9.0) under two conditions: with 5.3 M NaCl for halophilic activity assessment, or without NaCl for non-halophilic activity measurement. The reaction mixtures were lyophilized in a Freeze-dryer and the PET hydrolysis products were subsequently reconstituted with a 3-fold methanol solution for subsequent analysis.

### HPLC analytical procedure

HPLC analysis of PET hydrolysis products was performed using an Agilent 1220 system equipped with a ZORBAX SB-C18 column (4.6 × 250 mm, 5 μm, Agilent, Santa Clara, CA, USA). The system configuration included a pump module, an autosampler, a temperature-controlled column compartment maintained at 27 °C, and a UV detector set at 254 nm. The separation was achieved using a binary mobile phase system consisting of 0.1% TFA in water (solvent A) and 0.1% TFA in methanol (solvent B) at a constant flow rate of 1 mL/min. The gradient elution program was established as follows: 15% B (0-5 min), linear increase to 100% B (5-20 min), linear decrease to 15% B (20-25 min), and re-equilibration at 15% B (25-30 min). Quantification of TPA, MHET, and BHET was accomplished using external standard calibration curves.

### Transformer model for potential substitutions prediction

#### Training datasets collection

Homologous sequences of dsPETase05 were searched from Uniclust30 (version 2021_03) and the BFD database with HHblits (the number of iterations was set as 4, and other parameters were left as default values) using 17 thermophilic enzymes and 3 halophilic enzymes as seed sequences for queries (Supplementary Tables 6 and 7). All retrieved sequences were clustered using CD-HIT at 90% sequence identity, yielding 30,729 thermophilic homologous sequences and 16,959 halophilic homologous sequences.

#### Prediction of mutation sites

We applied Transformer model training as previously reported^13^. The models were trained for 20, 30, 50, and 100 epochs respectively using a batch size of 32. Residues were filtered based on the logits assigned to the wild-type amino acids. For each model, the top ten residue positions with the highest average prediction scores were selected. After removing duplicate amino acid positions, 45 wild-type residues exhibited lower fitness scores than potential substitution positions. (Supplementary Table 8).

### Construction of saturation mutants

The saturated mutant library was constructed using the NNK strategy^36^. Site-saturation mutagenesis was performed at 45 predicted positions by designing NNK primers (Supplementary Table 5) with dsPETase05 as a template. First, the plasmid pET28a-dsPETase05 was amplified by PCR using the designed NNK primers. The ligation products were then transformed into *E. coli* Rosetta gami 2(DE3) competent cells and plated on LB agar medium (Kan and Tc resistance), followed by overnight incubation at 37 °C. For each mutation site, 96 transformants were picked from the agar plates into a 96-well deep-well plate containing 400 μL LB liquid medium (Kan and Tc resistance) per well. The plate was sealed and incubated overnight at 37°C with 800 rpm shaking. Subsequently, 400 μL of 40% glycerol was added to each well, and the cultures were mixed and stored at −80 °C for later use. Sequencing validation and mutation coverage analysis were performed for two representative sites (P47 and F53) (Supplementary Fig. 7).

### High-throughput fluorescence screening

#### Strain activation

Frozen stock cultures were thawed on ice. Each well of a 96-deep-well plate was filled with 400 μL LB liquid medium (Kan and Tc resistance), and 10 μL of thawed culture was inoculated into each well. The plate was incubated overnight at 37°C with 800 rpm shaking for strain activation.

#### Protein expression

A fresh 96-deep-well plate was prepared with 400 μL LB liquid medium (Kan and Tc resistance) per well. Each well was inoculated with 25 μL of the overnight-activated culture and incubated at 37 °C with 800 rpm shaking for 3.5 h. The temperature was then reduced to 18 °C, and shaking continued for 0.5 h. Protein expression was induced by adding IPTG to a final concentration of 0.5 mM. After 24 h, cells were harvested by centrifugation at 4 °C and 4,500 *g* for 15 min. The supernatant was discarded, and cell pellets were flash-frozen in liquid nitrogen and stored at −80°C.

#### Cell lysis

A lysis buffer (0.5× BugBuster in 100 mM sodium phosphate buffer, pH 8.0) was prepared. The frozen cell pellets in the 96-well plate were thawed at 10 °C, and 300 μL of lysis buffer was added to each well. After mixing, the plate was incubated statically at 30 °C for 30 min. Cell debris was removed by centrifugation at 4 °C and 4,500 *g* for 10 min, and the clear lysate supernatant was collected for enzymatic assays.

#### Enzyme activity assay

FDL-PET films (1.6 mg/well, 0.1235 μg/mg PET) were prepared as substrates in black 96-well polypropylene plates according to the method described previously^12^. Reaction buffer (Tris-HCl, pH 9.0, 5.3 M NaCl) was added to each well (190 μL/well), followed by 10 μL of clear cell lysate. The mixture was gently pipetted to homogenize, sealed, and incubated for 24 h at the designated temperature. Fluorescence intensity (FI494/525) was measured using a microplate reader with excitation/emission at 494/525 nm. The enzymatic reaction mechanism and results are shown in Supplementary Figs 8 and 9, respectively.

### Determination of protein melting temperatures

To determine protein’s melting temperatures, a fluorescence-based thermal stability assay was employed. The SYPRO Orange dye 5000× stock solution in DMSO (Sigma-Aldrich, Germany) was first diluted to 250× in water. Protein samples were loaded onto a thin-walled 96-well PCR plate with each well containing a final volume of 25 μl. The final concentration of protein and SYPRO Orange dye in each well was 5 μM and 10×, respectively. The PCR plates were then sealed with optical quality sealing tape and spun at 3,000 rpm for 1 min at room temperature. The mixture was heated in a LC480 II real-time polymerase chain reaction (PCR) system (Roche, Switzerland) from 20 to 99 °C at a heating rate of 0.06 °C/s. Fluorescence measurements were conducted ten times per degree Celsius. The *T*_m_ was determined from the peak(s) of the first derivatives of the melting curve using the Thermal Shift Assay software.

## Data availability

Source data are provided with this paper. The atomic coordinates and structure factors of the reported structures have been deposited in the Protein Data Bank under accession codes 9VR5 for dsPETase01_apo, 9VR6 for dsPETase05_apo, 9VR7 for dsPETase05_TPA, 9VR8 for dsPETase05_S168A_MHET, and 9VR9 for dsPETase06_C23A_apo.

## Acknowledgements

This work was financially supported by the National Key Research and Development Project of China (grant no. 2022YFC2804500); National Natural Science Foundation of China (grant nos. 32370124 and 32025001); Shandong Provincial Natural Science Foundation (grant no. ZR2023ZD50); and Guangdong Basic and Applied Basic Research Foundation (grant no. 2025B1515020038).

## Author contributions

K.L., R.G., J.W., and S.L. conceived the research. G.Z., W.X., C.Zh., S.Z., J.D., N.W., X.C., G.L., Y.Z., and T.W. performed most of the bioinformatics analyses and mutation experiments. X.L., M.Z., J.H., C.C., S.H., and C.Z. determined all the protein structures. G.Z., X.L., and K.L. analyzed the results. Y.P., J.C., G.F. and X.Z. provided comments and suggestions on the paper. The paper was written and revised jointly by all the authors.

